# Translocational unfolding in clostridial binary iota toxin complex

**DOI:** 10.1101/721969

**Authors:** Tomohiro Yamada, Toru Yoshida, Akihiro Kawamoto, Kaoru Mitsuoka, Kenji Iwasaki, Hideaki Tsuge

**Affiliations:** Faculty of Life Sciences, Kyoto Sangyo University, Kamigamo-motoyama, Kita-ku, Kyoto, Japan; Institute for Protein Dynamics, Kyoto Sangyo University, Kamigamo-motoyama, Kita-ku, Kyoto, Japan; Institute for Protein Research, Osaka University, Suita, Osaka, 565-0871, Japan; Research Center for Ultra-High Voltage Electron Microscopy, Osaka University, Osaka, Japan; Life Science Center for Survival Dynamics, Tsukuba Advanced Research Alliance (TARA), University of Tsukuba, Tsukuba, Japan; Center for Molecular Research in Infectious Diseases, Kyoto Sangyo University, Kamigamo-motoyama, Kita-ku, Kyoto, Japan

## Abstract

Protein translocation across the membrane is critical for microbial pathogenesis and various cellular functions. Bacterial binary toxins such as anthrax toxin are composed of enzyme components and a translocation channel, which catalyses substrate unfolding and translocation. Here we report the structures of the clostridial binary toxin (iota toxin) translocation channel Ib-pore and its complex with ADP-ribosyltransferase Ia. The Ib-pore structure at atomic resolution provides a similar structural framework as observed for the catalytic ϕ-clamp of the anthrax protective antigen pore. However, the Ia-bound Ib-pore structure showed a unique binding mode of Ia: one Ia binds to the Ib-pore, and the Ia N-terminal domain interacts with Ib via two other Ib-pore bottlenecks with multiple weak interactions. Furthermore, Ib-binding induces Ia N-terminal α-helix tilting and partial unfolding, whereupon the unfolded N-terminus continues to the ϕ-clamp gate. This study reveals the novel mechanism of N-terminal unfolding, which is crucial for protein translocation.

## Introduction

*Clostridium perfringens* iota toxin (iota), *C. difficile* toxin(CDT), *C. spiroforme* toxin (CST), and *C. botulinum* C2 toxin belong to the family of binary toxins. It consists of the enzymatic ‘A’ component with actin-specific ADP-ribosyltransferase and the ‘B’ component which binds to the host cell and functions as the translocation channel of each enzymatic component (Ia,Ib; CDTa,CDTb; CSTa,CSTb; and C2I,C2II, respectively). The ‘B’ component precursor is first cleaved off by a cellular protease, then binds to the target cell via a receptor, forms a soluble oligomer termed the ‘prepore’, and finally converts to a membrane-spanning pore in an acidified endosome. The oligomer-receptor complex acts as a substrate docking platform that subsequently translocates an enzymatic component into the cytosol from the acidified endosome. Therein, the enzymatic component mono-ADP-ribosylates G-actin, inducing cytoskeletal disarray and cell death. The enzymatic component consists of two domains: the C-terminal domain with an actin-specific ADP-ribosyltransferase activity, and the N-terminal domain, considered as a binding domain to the membrane-spanning translocation component. The clostridial binary (iota) toxin consists of Ia and Ib^1–3^. However, although extensive structural and functional studies of Ia have been conducted^4,5^, little is known regarding the Ib translocation channel.

A similar binary toxin, anthrax toxin, constitutes a major virulence factor of *Bacillus anthracis*, consisting of an enzymatic component (two enzymatic proteins, edema (EF) and lethal (LF) factors) and a protein translocation channel (protective antigen (PA))^6^. LF is a zinc-dependent protease that cleaves mitogen-activated protein kinase kinase; the combination of PA and LF leads to the death of humans and animals. Cleavage of a PA precursor triggers heptamerization or octamerization (prepore), and the low pH endosomal environment causes the oligomer to insert into the membrane by forming a transmembrane β-barrel (pore)^7^. PA heptameric prepore structure was revealed with and without anthrax toxin receptor^8,9^. Furthermore, the LF-bound PA prepore complex structure (PA dimer with one LF N-terminal domain (LF_N_)) was revealed by crystallography at 3.1 Å resolution. The biological unit structure (PA_8_(LF_N_)_4_) was also deduced from the complex, revealing that the first N-terminal helix of LF binds on the PA dimer interface surface^10^.

Although PA pore instability has hampered its structural analysis, Jiang et al. clarified the PA heptameric pore structure by cryo-electron microscopy (cryo-EM) at 2.9 Å resolution^11^, revealing the catalytic ϕ-clamp and a long membrane-spanning channel^12^. The ϕ-clamp is the narrowest passageway of the PA translocation channel, in which the seven phenylalanine-427 residues converge within the lumen, generating a radially symmetric solvent-exposed aromatic ring. Krantz et al. proposed that the ϕ-clamp serves a chaperone-like function, interacting with hydrophobic sequences presented by the protein substrate as it unfolds during translocation^12^. Because translocation is driven by a transmembrane proton gradient, a Brownian ratchet model has been proposed, which depends on protonation and deprotonation of the translocating polypeptide acidic residues^12–15^. Furthermore, an allosteric helix-compression model regulated by the α-clamp in PA was recently proposed^16^, although it remains controversial whether this model is adequate.

To our knowledge, no high-resolution structure of the enzyme-bound translocation channel including the LF-bound PA-pore is available (although the PA heptamer pore structure with three LF_N_ was reported at 17 Å resolution^17^), and pore structural information remains limited in PA^11^ and other type Tc toxins^18,19^. In clostridial binary toxin, membrane prepore and pore structures have not been described. To understand the protein translocation mechanism via the bacterial translocation channel, here we determined the structures of the Ib heptameric translocation channel without and with the enzymatic component Ia (Ib-pore and Ia-bound Ib-pore, respectively) by cryo-EM at atomic resolution, providing the novel structural pose of Ia just prior to translocation in the transmembrane spanning Ib-pore.

## Results

### Ib oligomerization

Ib oligomerization is rapidly induced at 37°C in Vero cells in a temperature but not pH dependent-manner^20,21^. Conversely, *in vitro*, Ib oligomerizes poorly at 37°C and/or under acidic conditions following cleavage of the 20 kDa N-terminal propeptide. PA can oligomerize and form the PA-prepore upon propeptide cleavage, with the conformational change from prepore to pore being induced at low pH *in vitro*^*22*^; however, this leads to rapid and irreversible aggregation^11,23^. During cryo-EM sample preparation *in vitro*, we found that 10% ethanol could efficiently induce oligomerization upon N-terminal propeptide cleavage (Extended Data Fig. 1). Therefore, the sample was prepared by adding 10% ethanol and 0.03% lauryl maltose neopentyl glycol (LMNG) (Anatrace) following propeptide cleavage. Consequently, EM imaging showed that it led to the direct conversion of monomeric Ib to the pore, not the prepore. The obtained Ib-pore was stable at neutral pH, allowing cryo-EM structural analysis of the Ib-pore along with the Ia-bound Ib-pore.

### Ib-pore and Ia-bound Ib-pore cryo-EM data

Two data sets were collected using Titan Krios (FEI). With the first, we revealed the Ib-pore structure using 38,433 particles after classification of 299,491 extracted particles; local resolution analysis by Resmap^24^ showed the inner pore region at approximately 2.5 Å resolution (Fig. 1a, Extended Table 1, Extended Data Fig. 2). As described in Online Methods, the first data also included small amounts of Ia-bound Ib-pore particles, which provided a 5.2 Å resolution map of the Ia-bound Ib-pore (Extended Data Fig. 2).

**Figure 1.**
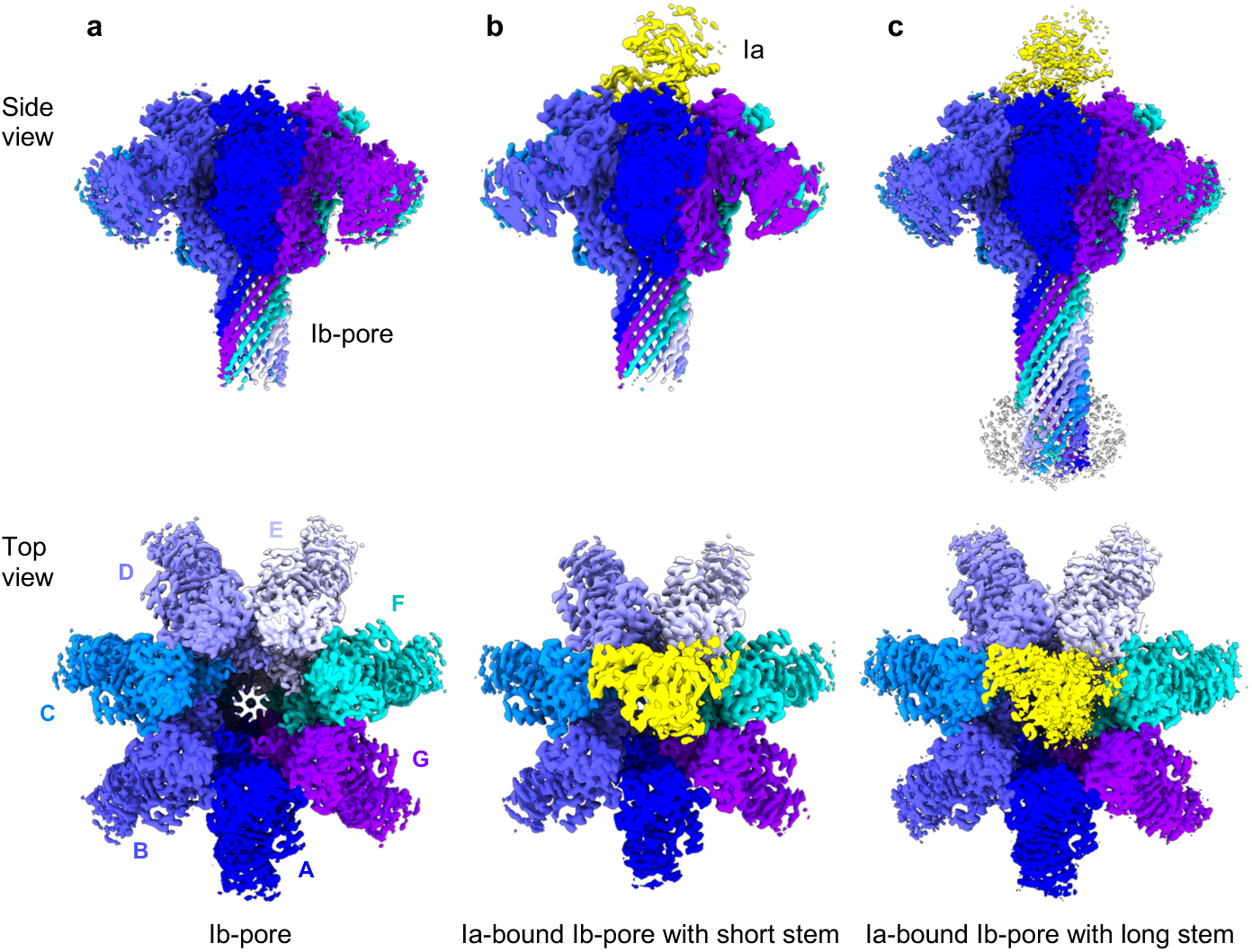
Cryo-EM density maps. **a,** Ib-pore with short stem. **b,** Ia-bound Ib-pore with short stem. **c,** Ia-bound Ib-pore with long stem. Ib protomers are shown in different colours.

Therefore, we generated another data set, raising the ratio of Ia. Two classes clearly showed Ia density on the Ib-pore with long β-barrel stem and short β-barrel stem, respectively. Therefore, using 135,359 particles for short stem class and 62,940 particles for long stem class following classification of 871,264 extracted Ia-bound Ib-pore particles, 3D refinement was performed after several additional classifications (Extended Data Fig. 3). Finally, 2.8 Å map (Ia-bound Ib-pore with short stem) and 2.9 Å map (Ia-bound Ib-pore with long stem) were yielded.

### Structure of the Ib-pore

The obtained cryo-EM map shows the Ib-pore having a funnel-like structure, lacking the lower stem region (328–365), suggesting that the bottom half of the stem had not yet formed. Thus, the Ib-pore structure contains a stem of 40 Å length (90 Å at full length, as noted for the Ia-bound Ib-pore), with the stem diameter the same as that of PA (>15 Å) (Fig. 2a-c; Extended Data Fig. 4a, 5a). The Ib-pore consists of four domains, 1′ (domain 1 without the propeptide), 2, 3, and 4, as with PA. The main pore body comprises domain 2, which consists of two parts designated as 2c (residues 296–311 and 381–512) and 2s (residues 312–380) (Fig. 2a, c). Domain 2s is an extended β-hairpin, seven copies of which assemble to form a membrane-spanning 14-stranded β-barrel. Domain 3 is located at an intermediate position between domains 1′ and 2c. The cryo-EM density of domain 4, the receptor-binding domain, is weak and has the lowest resolution among all four domains (Extended Data Fig. 2), likely owing to its flexibility resulting from minimal contact with the other domains. The overall structure of the Ib-pore is similar to that of the PA-pore (Extended Data Fig. 5a). Specifically, the funnel structure consisting of domains 1′, 2, and 3 shows high similarity with that of the PA-pore. Domains 1 and 2 of PA share 41 and 40% sequence identity with the corresponding regions of Ib (Extended Data Fig. 6). The weak map density of domain 4 is also reported for the PA-pore^11^. In Ib, the relative position of this density differs from that of PA-pore domain 4 (Extended Data Fig. 5a). As domain 4 is a receptor binding domain, the sequence identity shows < 10% identity with PA, with the size also differing (Ib domain 4 is twice as large as that of PA).

**Figure 2.**
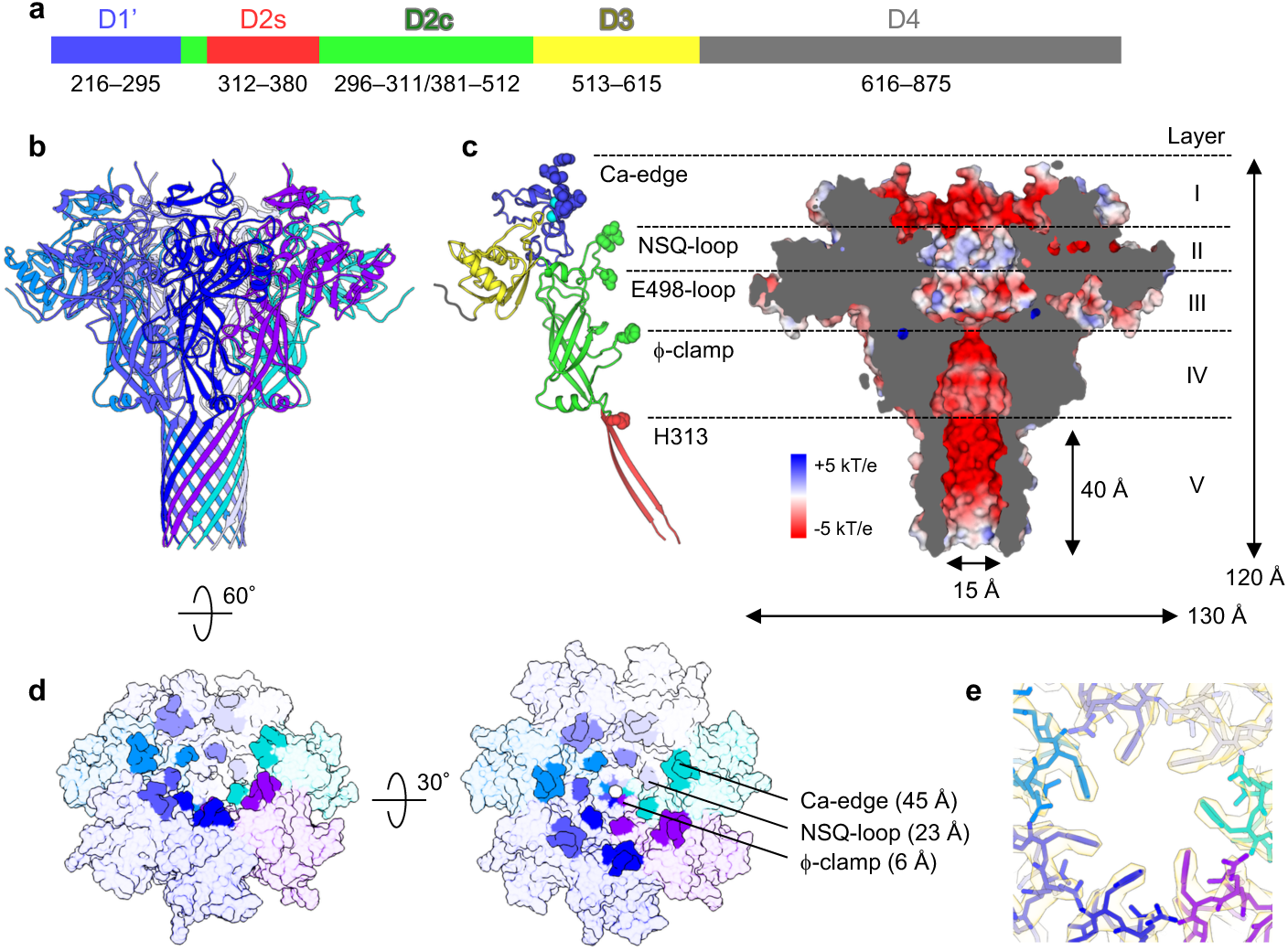
Atomic model of the Ib-pore. **a,** Ib Domain structure. **b,** Overall structure of the Ib-pore with short stem. The domain IV model was not built. **c,** Ib-pore layer structure. Five layers are shown in a protomer cartoon model colour-coded as in **a** along with the cut-away surface electrostatic potential at pH 7.0. The amino acid residues forming the boundary between adjacent layers are shown as a sphere model. **d,** Three bottlenecks of the Ib-pore. Ca-edge (216–224), NSQ-loop (490–492), and ϕ-clamp (454). Surface model is shown. Diameters of bottlenecks are shown in parentheses. **e,** Top view of the f-clamp with cryo-EM density map.

The cryo-EM density of the inner side of the funnel was analysed at high resolution; thus, the side chain is clearly visible (Extended Data Fig. 4b). The narrowest clamp is formed by seven F454s from seven protomers with a diameter of 6 Å, termed the ϕ-clamp in PA (Fig. 2e). Two additional bottlenecks exist in the cis-side: Ca-edge (216–224), an N-terminal Ca binding site, and NSQ-loop (490–492), with 45 Å and 23 Å diameters, respectively (Fig. 2d). Ca-edge, which is a unique di-calcium binding site (Extended Data Fig. 5b), and NSQ-loop are the most important regions for Ia-binding as described later. We then divided the inner pore surface as layer I, II, III, IV, and V from the cis- to trans-side (Fig. 2c). Layer I represents the broadest area of the funnel from the Ca-edge to NSQ-loop. Layer II is from the NSQ-loop to E498-loop. Layer III is from the E498-loop to ϕ-clamp. Layer IV is from the ϕ-clamp to H313 in the trans-side. We designated the stem region as layer V. The long stem with a β-barrel is created by an amphipathic flexible loop (E312–I380) of the prepore.

The enzymatic component Ia (ADP-ribosyltransferase) is translocated from the cis (layer I, II, and III) to the trans-side (layer IV and V). Although the sequence is not well-conserved on the inner surface between the Ib-pore and PA-pore, structural similarities exist and the negative (pH 7) and positive (pH 5.5) surface potential in the cis side (Layer II and III) are maintained in both channels (Extended Data Fig. 5c). In PA, N422 and D425 (422–NAQDDFSST–430) were noted as constituting an essential pH sensor to lead the conversion from prepore to pore; in Ib, these two residues are conserved. Notably, H313 locates on the inner surface of the upper stem, which appears to have an important function for translocation (Fig. 2c). Moreover, numerous Ser and Thr residues exist in the inner surface stem region, which may have vital functions as with PA although the residue positions are not conserved between Ib and PA (stem region Ser/Thr content of 56% each in Ib and Pa) (Extended Data Fig. 5c).

### Structure of the Ia-bound Ib-pore

Two separated classes, Ia-bound Ib-pore with short and long stem, were obtained from the second data set by C1 data analysis (Extended Data Fig. 3). In both structures, Ia sits on the cis-side of the Ib-pore with the same binding mode (Fig. 1b-c and 3a-c; Extended Data Fig. 4a). Notably, the Ia stoichiometry and binding mode differed from those of the LF-bound PA-pore (PA:LF = 8:4 or 7:3). Clearly, one Ia binds to the Ib-pore heptamer. Half of the Ia N-terminal domain is buried in the Ib-pore through several interactions with interface 1650 Å^2^. They primarily interact asymmetrically with Ca-edges and NSQ-loops. Specifically, five Ca-edges in subunit C-G contribute to Ia binding, with seven NSQ-loops in subunit A-G supporting the Ia N-terminal domain (Fig. 3d-g; Extended Data Fig. 7).

**Figure 3.**
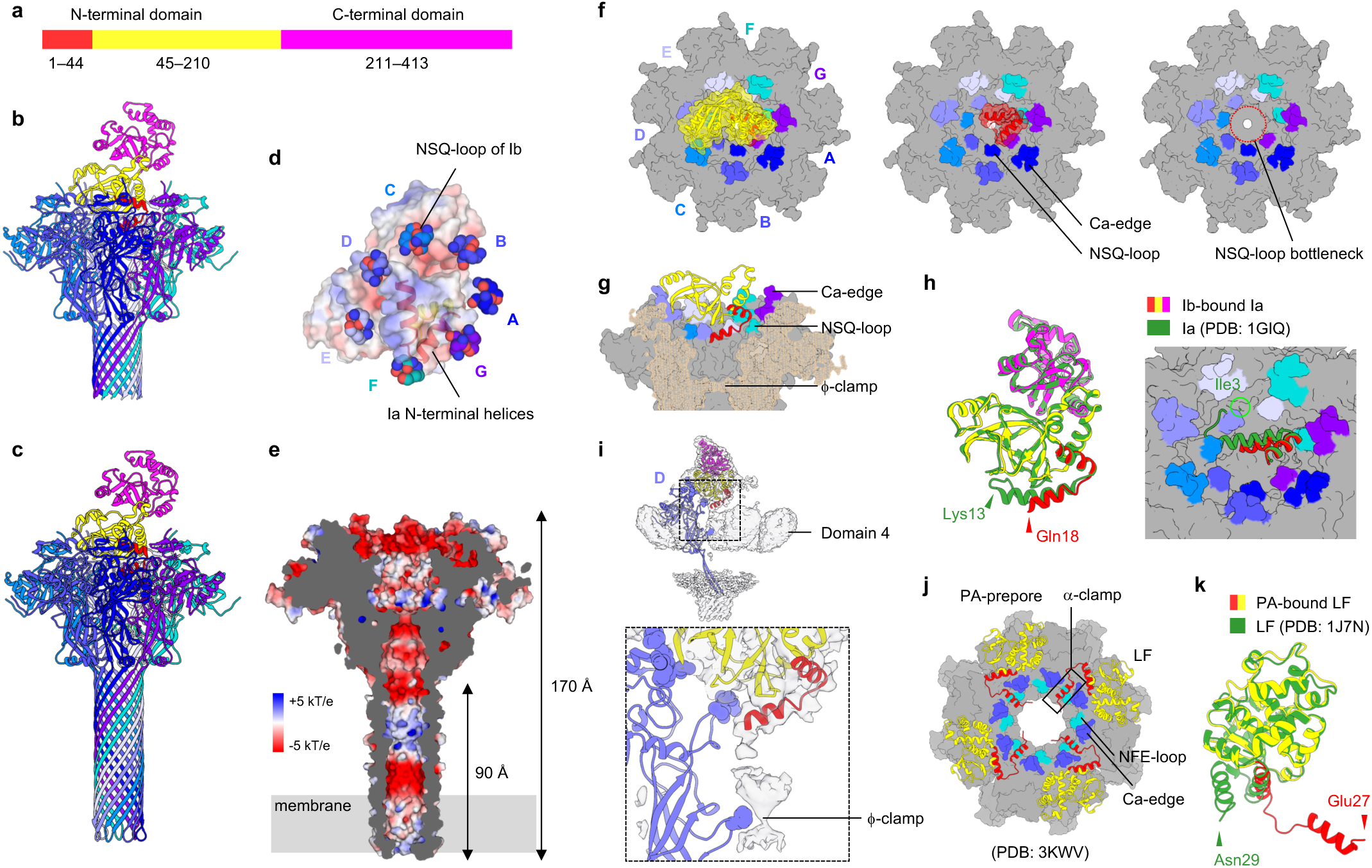
Atomic model of the Ia-bound Ib-pore. **a,** Ia domain structure. **b,c** Ia-bound Ib-pore overall structures with short and long stems. **d,** Ia, bottom view. **e,** Cut-away surface electrostatic potential at pH 7.0. **f,** Top views. For clarity, 1–210 (left) and 1–44 (center) regions of Ia are shown. Ia is not shown (right). **g,** Cut-away view. **h,** Unfolding of Ia N-terminus. Ib-bound Ia and Ia crystal structure are superimposed. **i,** Extra map leading from the N-terminal helix to ϕ-clamp. Map within 3 Å of the Ib model is subtracted from the original map. **j,** LF-bound PA-prepore structure. **k,** PA-bound LF and LF crystal structure superposition.

No large Ia-induced structural change occurs in the Ib-pore between the apo-Ib-pore and Ia-bound Ib-pore; only two NSQ-loops (E and F subunits) exhibit asymmetric positional change (Fig. 3f). Moreover, no large structural change occurs in almost the entire Ia molecule except the N-terminal (1–44) region between apo-Ia and the Ia-bound Ib-pore (Fig. 3h). Within the seven NSQ-loop interaction, one (F subunit) pushes residues 29–32 in the N-terminal α-helices (Ia), causing the apparent tilt and partial unfolding of the N-terminal α-helix (Fig. 3h; Extended Data Fig. 7). Thus, the N-terminal region (1–17) in Ia unfolds in the Ib-pore because it is too large to fit in the NSQ-loop bottleneck, then continues to the Ib-pore ϕ-clamp gate (Fig. 3h, i). It appears that the Ia molecule floats from the Ib protomers via NSQ-loops, thus providing free inner space to accommodate the Ia N-terminal region. In summary, the main interactions are caused by several asymmetrical Ca-edges and NSQ-loops. No specific strong interaction exists between Ia and the Ib-pore, suggesting that this may constitute an essential feature for the translocation channel and substrate protein, with the weak interactions affording efficient translocation.

Although detailed reports regarding Ia–Ib binding are lacking, an earlier study showed that the N-terminal domain (residue 216–321 in domain 1′) of Ib is essential for Ia docking^25^. This cryo-EM study of the Ia-bound Ib-pore provides precise information regarding Ia and Ib-pore interactions, showing the Ca-edge of domain 1′ along with the NSQ-loop of domain 2 as essential for binding. It was also reported that Ib lacking just the first N-terminal 27 residues did not facilitate Ia entry^25^. This is because the N-terminal 27 residues form the Ib-pore Ca^2+^ binding site (Ca-edge) (Extended Data Fig. 5b). Moreover, residues 129–257 were proposed as the minimal Ia fragment for translocation^26^. The present Ia-bound Ib-structure showed that a native ordered structure of the Ia N-terminal domain (45–210) is necessary for the stacking via NSQ-loops.

We next compared previous mutational results of Ib with the present Ib-pore structure. F454A led to loss of cytotoxicity and markedly increased single-channel conductance, suggesting that the ϕ-clamp is highly conserved and crucial for binary toxin activity^27^. Several mutations within the amphipathic β-strand forming the stem affected pore formation, single-channel conductance, and ion selectivity (S339E–S341E, Q345H, and N346E)^27^. Based on the structure, S339, S341, and N346 are located on the inner stem surface, whereas Q345 is found on the stem tip. Ser and Thr residues on the inner surface are likely essential for translocation.

## Discussion

The available cryo-EM structure of LF_N_-bound PA-pore (PA_7_(LF_N_)_3_) is at low resolution; nevertheless, the same binding mode as in the LF_N_-bound PA-prepore is assumed^10,17^. The first α-helix and β-strand of LF_N_ unfold and dock into the deep amphipathic cleft on the octamer surface, termed the ‘α-clamp’ (Fig. 3j). Thus, the main interactions are formed by the N-terminal helix binding to the PA-prepore α-clamp. Altough similar structural feature and electrostatic potential exist for Ib-pore translocation channel, significant differences exist between Ia and LF binding. For LF, the N-terminal α-helix binds to the PA-pore α-clamp, with the following 26 N-terminal residues being invisible owing to their flexibility. This flexibility is also observed in the apo-structure, suggesting that the N-terminal region is intrinsically flexible (Fig. 3k). Conversely, for Ia, binding to the channel causes Ia N-terminal unfolding. This indicates that the Ib-pore serves as an unfolding chaperone for substrate translocation even at neutral pH. As the Ib α-clamp site does not function as an α-clamp, the Ib-pore rather uses a novel mechanism for Ia N-terminal region unfolding: (1) a large portion of the N-terminal domain of the Ia structure lies in the Ib-pore, and (2) the interaction induces Ia N-terminal helix tilting and partial unfolding (Fig. 3h). Thus, the Ib-induced disordered region (1–AFIERPEDFLKD–12) followed by unfolded N-terminal helix (13–KENAI–17) directly continues to the ϕ-clamp (Fig. 3i).

The Ia N-terminal region contains numerous hydrophobic along with both positive and negative residues. Ia and LF share no sequence similarity, suggesting that the characteristic residue assortment (positive, negative, and hydrophobic) is essential for translocation. Constructs lacking both negative and positive charges in the unstructured region of LF_N_, composed of only Gly, Ser, and Thr, translocate more slowly and independently of the ΔpH, indicating that a balance of acidic- and basic-charged residues is required for efficient translocation with ΔpH^28^. Furthermore, in endosomes, as the phenylalanine clamp could be considered as a barrier of pH difference, the cis-side (Layer II and III) electrostatic potential is positive (pH 5.5), whereas the trans-side (IV) electrostatic potential is negative (pH 7.0) (Extended Data Fig. 5c). The electrostatic potential difference represents a common aspect in Ib-pore and PA-pore and is likely important for translocation. It is also noted that electrostatic repulsion between pore (NSQ-loop in subunit E) and substrate protein (Arg26) seems to be important for efficient translocation (Extended Data Fig. 7b). Furthermore we consider that the destabilization of Ca-edge at endosomal acidic pH is also key to reduce the interaction, leading to more efficient translocation.

Despite the differences of binding stoichiometry and binding mode of Ia/Ib and LF/PA, the unfolded Ia and LF N-terminals are accommodated in electrostatically charged cis-side pore, then led to the ϕ-clamp. Therefore, we consider that the first entry event, in which the tip of the unfolded N-terminus enters into the gate of the hydrophobic ϕ-clamp, is significant for translocation, similar to threading through a needle. For Ia, the unfolded N-terminal region becomes freely accessible to the ϕ-clamp in the space of Layer III under the NSQ-loop. Thus, Ia N-terminal movement in the limited space is beneficial compared with LF-bound PA with large open space, allowing the hydrophobic tip to readily reach the hydrophobic ϕ-clamp gate. This first event appears necessary for an extended-chain Brownian ratchet model.

Two ϕ-clamp configuration states (clamped and unclamped dilated states) have been proposed that are allosterically regulated by the α-clamp^16,29^. The allosteric helix-compression model was proposed as more favourable than the extended-chain Brownian ratchet model. This model explains that the successive α-helix formation induced by the α-clamp is essential for substrate translocation and that newly formed α-helices pass through the dilated ϕ-clamp, leading to produce more power stroke. However, structures of both the PA-pore at acidic pH and Ib-pore at neutral pH show that the ϕ-clamp forms the same configuration as in the closed (clamped empty) state. The dilated (unclamped empty) state structure has not yet known; moreover, whether the dilated state exists and the allosteric helix-compression model is generalizable remain controversial^30,31^. In Ib and C2II, the dilated state has not been observed by electrophysiological study^32,33^ or in the present cryo-EM studies. With regard to its structure, as seven phenylalanines are stacked in the ϕ-clamp structure, change to a more dilated conformation is unlikely. In addition to these, Ib does’t use the α-clamp for α-helices binding. Together, these observations indicate that the Ib translocation likely occurs via a static ϕ-clamp pore, suggesting the extended-chain translocation of the unfolded N-terminal substrate (Fig. 4). The presented binary toxin complex structure and the mechanism of unfolded N-terminal substrate translocation should be conserved in other *C. difficile, C. spiroforme*, and *C. botulinum* binary toxins. Notably, our study provides structural clues to develop inhibitors of these binary toxins, especially CDT from human opportunistic pathogen *C. difficile*^34–36^ or iota-like toxin (CPILE/BEC) in human food poisoning outbreaks^37–39^.

**Figure 4.**
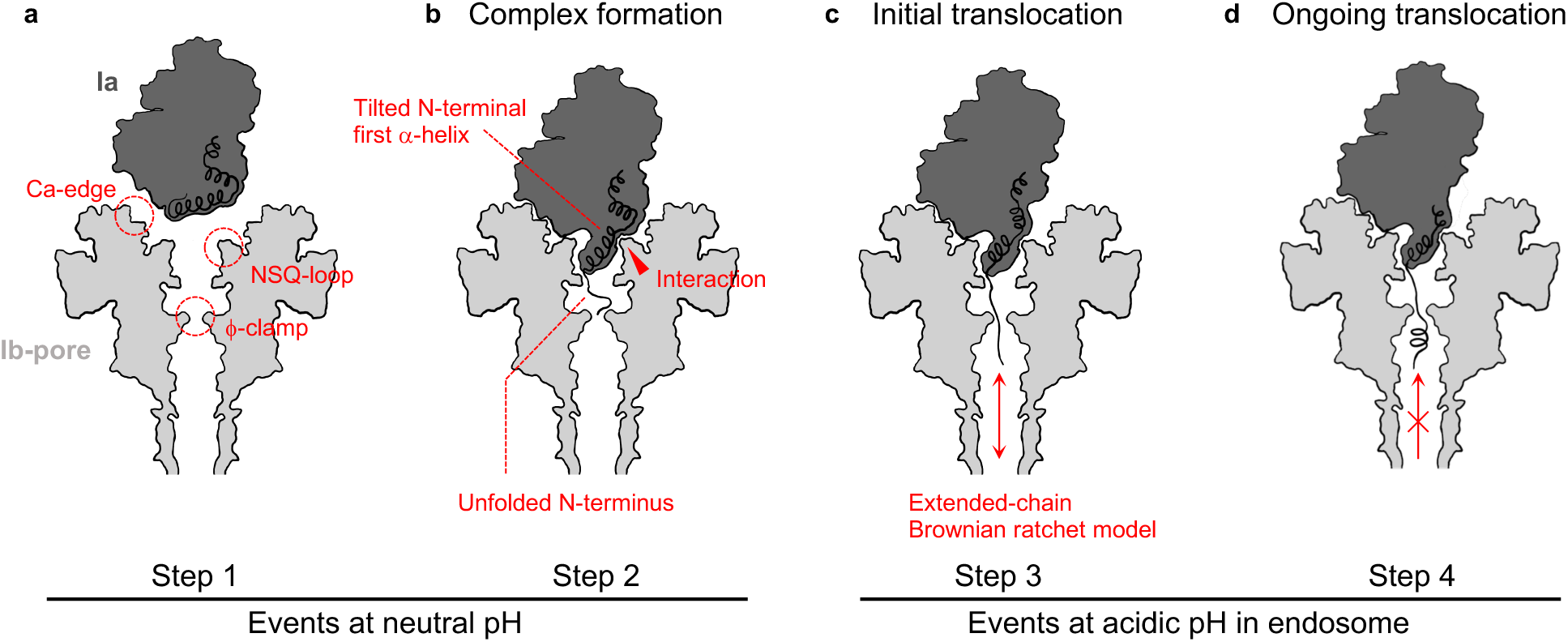
Translocation model of Ia via Ib-pore. Ia (black) and Ib-pore (grey). Arrows indicate possible Ia movement direction. **a,** Ia and Ib-pore before complexation. **b,** Ia-bound Ib-pore complex solved herein by single particle analysis. Ib-pore binding induces tilts and partial unfolding of the first N-terminal Ia α-helix. **c**, Endosomal acidic conditions facilitate unfolded N-terminal tip entry into the Ib-pore ϕ-clamp followed by extended-chain Brownian ratchet model-mediated translocation. **d,** Expected mechanism to prevent Ia retro-translocation by α-helix formation in the stem at neutral pH in the trans-side.

At neutral pH, only a minor conductance decrease was observed upon Ia addition to the membrane cis-side^32^. However, at pH 5.6, cis-side conductance decreased to 30–40% of the open configuration conductance^32^. Ia-mediated Ib blockage occurred only at pH 5.6, suggesting the translocation occurs only at acidic pH. In our study, we revealed the Ia-bound Ib-pore structure at neutral pH. The disordered Ia N-terminal region tip reaches the ϕ-clamp (Fig. 3i); however, it may not be entirely blocked but still fluctuate in the pore.

In summary, the present structure captured the Ia-bound Ib-pore just before translocation: The Ia N-terminal region exhibited Ib-induced unfolding. In future studies, using the stable Ib-pore, we expect to capture the translocation state of Ia in the Ib-pore under acidic conditions.

## Acknowledgements

This work was supported by JSPS KAKENHI Grant Numbers 18K06170 and 17K15095. This work was also supported by the Basis for Supporting Innovative Drug Discovery and Life Science Research (BINDS) from the AMED and the “Nanotechnology Platform” of the Ministry of Education, Culture, Sports. We thank H. Murata for the initial purification of Ib and JI. Kishikawa for help in cryo-EM analysis. HT thanks M. Nagahama and M. Oda for helpful comments on the studies.

## Author contributions

T.Yamada, T.Yoshida, A.K., and H.T. participated in research design and data analyses; T.Yamada prepared the Ib-pore and Ia-bound Ib-pore for cryoEM; T.Yamada, A.K., K.M., and K.I. performed cryoEM data acquisition and image processing; T.Yoshida performed the atomic model building, structure refinement and analyses; all author contributed to writing the manuscript and H.T. supervise the project.

## Author information

Reprints and permissions information is available at www.nature.com/reprints

### Competing interests

The authors declare no competing interests.

## Methods

### Ib and Ia expression and purification

The *iota toxin b* (*Ib*) gene was cloned into pGEX4T-1 without a signal peptide, and Ib was overexpressed in *Escherichia coli* Origami. The transformant was cultured in super broth medium containing ampicillin (50 μg/ml), tetracycline (12.5 μg/ml), and kanamycin (15 μg/ml) at 37°C until OD_600_ of 0.6, then isopropyl β-D-1-thiogalactopyranoside (final 1 mM) was added followed by culturing at 23°C for 16 h. The harvested cells were resuspended in lysis buffer containing 20 mM Tris pH 8.0, 150 mM NaCl, 2 mM CaCl_2_, and 5 mM dithiothreitol, and disrupted by sonication. After centrifugation at 180,000 × *g* for 40 min, the supernatant was loaded onto a Glutathione Sepharose 4B resin (GE Healthcare) column. After washing the column with lysis buffer, the bound protein was eluted by a buffer containing 20 mM Tris pH 8.0, 150 mM NaCl, and 10 mM reduced glutathione. The eluted fractions were concentrated to 19 mg/ml and the buffer was exchanged to 20 mM Tris pH 8.0, 50 mM NaCl, 2.5 mM CaCl_2_.

The *iota toxin a* (*Ia*) gene was cloned into pET-15b or pET-21b to produce Ia with N-terminal or C-terminal His-tag, respectively. Then, Ia was overexpressed in *E. coli* BL21 Star (DE3). The transformant was cultured in super broth medium containing ampicillin (50 μg/ml) at 37°C until OD_600_ became 1.5 and then isopropyl β-D-1-thiogalactopyranoside was added (final 0.5 mM) followed by culturing at 37°C for 16 h. The harvested cells producing each Ia were individually resuspended in lysis buffer containing 20 mM Tris pH 8.0, 300 mM NaCl, and 20 mM imidazole, and disrupted by sonication. After centrifugation at 180,000 × *g* for 40 min, the supernatant was loaded onto a Ni-NTA agarose column. After washing the column with lysis buffer, the bound protein was eluted by a buffer containing 20 mM Tris pH 8.0, 300 mM NaCl, and 500 mM imidazole. The eluted fractions were concentrated and the buffer exchanged to that containing 20 mM Tris pH 8.0, 2 mM CaCl_2_ to load onto an HiTrap Q HP 5 ml column (GE Healthcare). After anion exchange purification, Ia with C-terminal His tag was concentrated to 2.63 mg/ml, and the buffer was exchanged to 10 mM Tris pH 8.0 and 100 mM NaCl. Alternatively, Ia with N-terminal His-tag was incubated with 0.001 (unit/μg Ia) thrombin for 16 h at room temperature. Ia, in which the N-terminal His-tag was cleaved, was loaded onto a Ni-NTA agarose column, then the flow-through and wash fractions were collected. Collected fractions were concentrated to 22 mg/ml and buffer was exchanged to 10 mM Tris pH 8.0 and 100 mM NaCl using an Amicon filter.

### Sample preparation for the first data set

Ib oligomerizes poorly at neutral and acidic pH *in vitro* after cleavage of the pre-sequence with α-chymotrypsin. Ib (30 mg) was treated with 30 μg α-chymotrypsin for 1 h at room temperature. This reaction was terminated by adding phenylmethylsulphonyl fluoride (PMSF) (final 1 mM). Then, Ia with C-terminal His-tag was added to the Ib solution with three-fold molar excess and incubated 1 h at 37°C. The solution was loaded onto a Ni-NTA agarose column. We expected that the elution fraction includes Ia-bound Ib oligomer, but it was failed because they didn’t coexist in high concentration of imidazole. From oligomerization screening, we found that ethanol induced oligomerization of Ib efficiently after cleavage of pre-sequence incubating with α-chymotrypsin (Extended Data Fig. 1a). Accordingly, we changed to use the ethanol oligomerization to apply the flow-through fraction including Ib. Ethanol (final 10%) and LMNG (final 0.03%) was added to 1 mg Ib in the flow-through fraction and incubated at 37°C for 1 hour. In order to separate Ib oligomer from other small proteins, the solution was loaded onto 10.5 ml of Glycerol gradient bed (10–30% Glycerol, 50 mM HEPES pH 7.5, 100 mM NaCl, 1 mM CaCl_2,_ 0.03% LMNG). After ultracentrifuge at 230,139 × *g* for 16 hours, fractions were collected by 250 μl from bottom. Fractions showing high molecular mass by sodium dodecyl sulphate-polyacrylamide gel electrophoresis (SDS-PAGE) were collected and buffer was exchanged to 10 mM HEPES pH 7.5, 1 mM CaCl_2_, and 0.01% LMNG and concentrated to 1.9 mg/ml. This sample included Ib-pore and a small amount of Ia (Extended Data Fig. 1b, c).

### Sample preparation for the second data set

During sample preparation for the first data set, binding between Ia and the Ib-pore occurred and the LMNG concentration was too high for cryo-EM data collection. To solve these problems, Ib was purified with a smaller concentration of LMNG and Ia was added following Ib-pore purification. Ib (21 mg) was cleaved by 1 μg α-chymotrypsin for 1 h at room temperature. This reaction was terminated by adding PMSF (final 1 mM). The solution was then treated with 10% ethanol and 0.03% LMNG for 1 h at 37°C. To separate Ib oligomer from other small proteins, the solution was loaded onto 10.5 ml of Glycerol gradient bed (10–30% Glycerol, 50 mM HEPES pH 7.5, 100 mM NaCl, 1 mM CaCl_2,_ 0.003% LMNG). After ultracentrifugation at 230,139 × *g* for 16 h, 250 μl fractions were collected from the bottom. Fractions showing high molecular mass by SDS-PAGE were collected and the buffer was exchanged to 10 mM HEPES pH 7.5, 1 mM CaCl_2_, and 0.003% LMNG. Purified Ib oligomer was concentrated to 2.4 mg/ml, then Ia without His-tag was added with three-fold molar excess at the final step (Extended Data Fig. 1d, e).

### Cryo-EM imaging of the Ib-pore for the first data set

Sample vitrification was performed using a semi-automated vitrification device (Vitrobot Mark IV, Fisher Scientific, Eindhoven, The Netherlands). A 2.6 μl aliquot of sample solution at a concentration of 0.38 mg/ml (1/5 dilution of the sample) was applied to glow-discharged Quantifoil R1.2/1.3 in the Vitrobot at 100% humidity. The grid was then automatically blotted once from both sides with filter paper for 4.5 s blot time. The grid was then plunged into liquid ethane with no delay time. Cryo-EM imaging was performed using a Titan Krios (Fisher Scientific) operating at 300 kV acceleration voltage and equipped with a Cs corrector (CEOS, GmbH) and a direct electron detector Falcon 3 (counted mode) (Fisher Scientific) in automated data collection mode at a calibrated magnification of 1.13 Å/pixel (magnification ×59,000) and dose of 50 e/Å^2^ (or 0.46 e/Å^2^ per frame) with total 84.09 s exposure time. The data were automatically collected using EPU software with a defocus range of −0.8 to −2.5 μm and were fractionated 108 movie frames.

### Image processing of the Ib-pore for the first data set

A total of 2,120 images were collected in the first data set. The movie frames were subsequently aligned to correct for beam-induced movement and drift using MOTIONCORR2^40^, and contrast transfer function (CTF) were estimated using CTFFIND4^41^. A total of 299,491 particle images were automatically picked using Gautomatch (http://www.mrc-lmb.cam.ac.uk/kzhang/) and several rounds of 2D classification and 3D classification were performed using RELION-3.0^42^. The best among the 3D classes in which clearly showed 7-fold rotational symmetry in Ib-pore were subjected to 3D refinement with C7 symmetry. The 3D-refined structure was further CTF refined using the per-particle defocus and Bayesian polishing, which improved the resolution to 2.9 Å and a B-factor of −46 Å^2^ without the substrate Ia (Extended Data Fig. 2a-e).

### Image processing of the Ia-bound Ib-pore for the first data set

While the 3D class reconstruction proceeded in Ib-pore analysis, we identified another class that showed Ia density on the Ib-pore (1,735 particles). The class was subjected to 3D refinement and used as a template for a second 3D classification with 154,378 particles. Classes that exhibited density on the Ib pore were selected for processing using 3D refinement. Around the Ib pore membrane spanning stem (outside of the stem), some blurred density was observed that appeared irregularly in each class; therefore, it was subtracted to increase the efficiency of classification. Subtracted particles were subjected to a third 3D classification, and then classes that contained 15,890 particles with strong density on the Ib pore were subjected to 3D refinement initially without and then with a solvent mask. Finally, an Ia-bound Ib map was generated at 5.2 A□ resolution (Extended Data Fig. 2f-j). This result prompted us to collect a second data set using the sample while raising the ratio of Ia.

### Cryo-EM imaging of the Ia-bound Ib-pore for the second data set

A 2.6 μl aliquot of sample solution at a concentration of 0.48 mg/ml (1/5 dilution of the sample) was applied to glow-discharged Quantifoil R1.2/1.3. Other procedures including sample vitrification, blotting, freezing, and Cryo-EM imaging were the same as described for the first data set.

### Image processing of the Ia-bound Ib-pore for the second data set

Image processing was performed as described for the first data set unless otherwise stated. A total of 2,151 images were collected in the second data set. The movie frames were subsequently aligned to correct for beam-induced movement and drift using MOTIONCORR2^40^, and CTF were estimated using GCTF^41^. A total of 871,264 particle images were automatically picked using Gautomatch and several rounds of 2D classification were performed using RELION-3.0^42^. A total of 335,767 particles in the best class were subjected to 3D refinement, per-particle CTF refinement and Bayesian polishing. The polished particles were subjected to 3D classification using 3D refinement structure as the reference and divided into eight classes. The two classes clearly showed Ia density on the Ib-pore with long stem and short stem, respectively. These two classes were subjected to 3D refinement, per-particle CTF refinement and Bayesian polishing. The resulting 3D structure was further subjected to no-align 3D classification using a mask covering Ia. The final 3D refinement and postprocessing of two classes yielded maps with global resolution of 2.91 Å and B factor of −29 Å^2^ (Ia-bound Ib-pore with long stem) and 2.80 Å and B factor of −10 Å^2^ (Ia-bound Ib-pore with short stem), according to 0.143 criterion of the FSC (Extended Data Fig. 3).

### Model building and refinement

An Ib-pore model with short stem was built using the first data set. An initial rigid-body fit of PA structure (PDB ID: 3J9C) was applied into the cryo-EM density map using UCSF Chimera^43^. The Ib-pore model was then manually built by iterative rounds of model modification in COOT^44^ and refinement using PHENIX Real Space Refinement with secondary structure restraint^45^. Model building of Ib Domain 4 was not carried out because of the low resolution and lack of available crystal structure.

An Ia-bound Ib-pore model with short stem was built using the cryo-EM density map with partial stem from the second data set. Following initial model building and rigid-body fit using the Ib-pore model from the first data and the crystallographic structure of Ia (PDB ID: 1GIQ), they were manually modified and refined by iterative rounds of COOT and PHENIX as for model building of the Ib-pore with short stem.

Furthermore, an Ia-bound Ib-pore model with long stem was built using the cryo-EM density map with intact stem from the second data set. Following rigid-body fit of the Ib-pore model with short stem, the intact long stem was manually built using COOT and they were modified and refined by iterative rounds of COOT and PHENIX. Although the density of Ia (C-terminal domain) was insufficient for *de novo* model building, the cryo-EM density maps with short and long stem showed that the Ia and Ib conformations of the two maps were the same. Therefore, rigid-body fitting of the Ia model from the Ia-bound Ib-pore with long stem was finally carried out using UCSF Chimera. Then, the final Ia-bound Ib-pore with long stem model was refined using PHENIX Real Space Refinement with secondary structure restraint.

In the manuscript, we discussed Ia binding using the Ia-bound Ib-pore model with short stem because the cryo-EM density of Ia with short stem was clearer than that with long stem. The structures with long and short stem are the same except for stem length. The difference of cryo-EM density of Ia was likely attributed to the difference of particle number (short:135,359 and long: 62,940).

All figures were prepared using PyMOL (https://pymol.org/2/), UCSF Chimera, and UCSF ChimeraX^46^.

## Data availability

Cryo-EM maps and coordinates were deposited in the Electron Microscopy Data Bank and Protein Data Bank with the accession codes EMDB-0721 and PDB 6KLX for the Ib-pore, EMDB-0713 and PDB 6KLO for the Ia-bound Ib-pore with short stem, and EMDB-0720 and PDB 6KLW for the Ia-bound Ib-pore with long stem.

