## Extended Data for "Translocational unfolding in clostridial binary iota toxin complex"

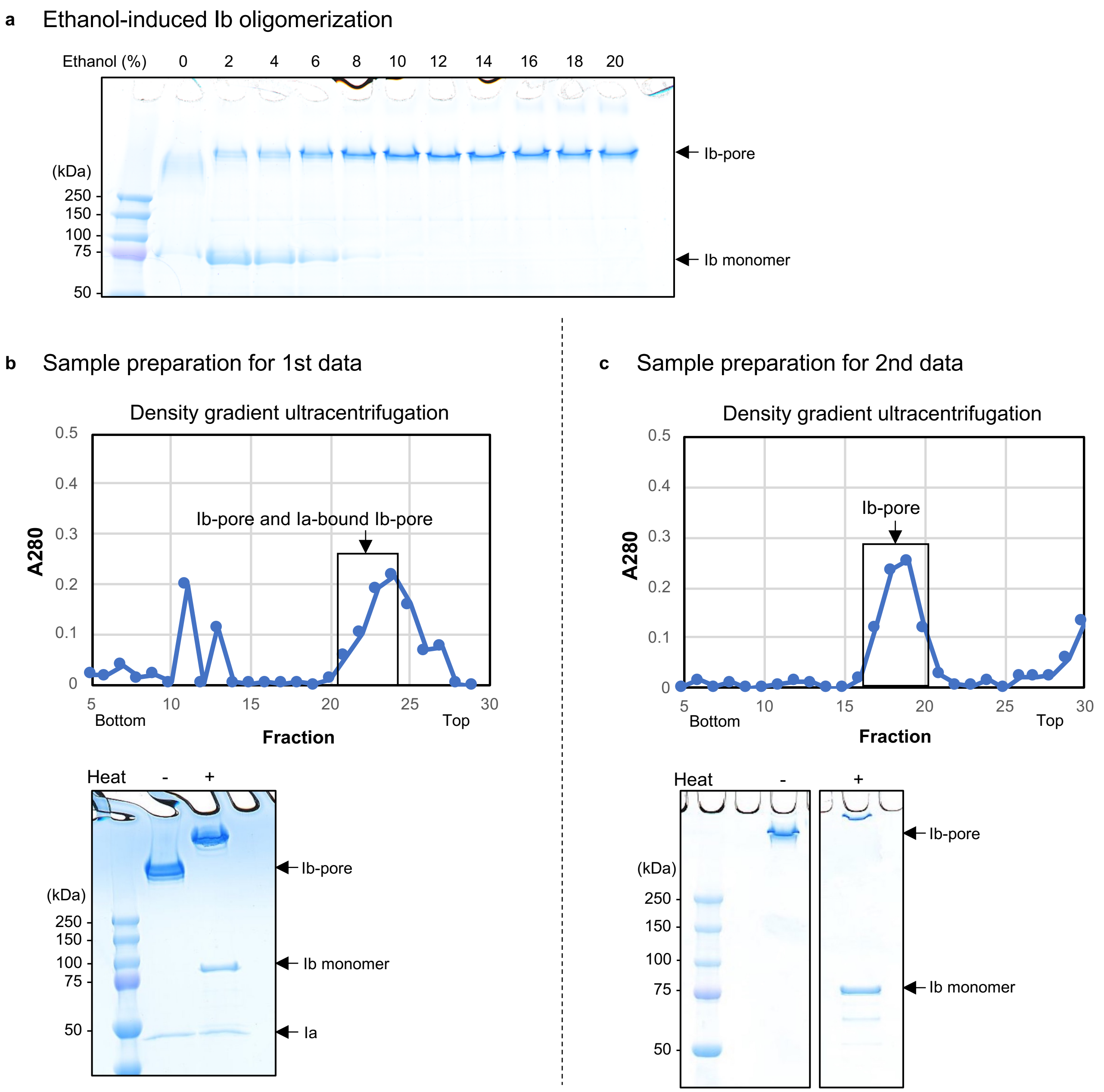


**Extended Data Figure 1 | Sample preparation of iota toxin.**

**a,** Ethanol-induced Ib oligomerization. After treatment with a-chymotrypsin, Ib monomer was oligomerized by adding ethanol. Oligomerization efficiency reached almost 100% in the presence of 10% ethanol. High concentration of ethanol (around 20%) caused aggregation, reflected in the band above the Ib-oligomer. **b,** Sample preparation for 1st data. *top*, Density gradient ultracentrifugation. *bottom*, SDS-PAGE of fractions 21–24, showing that these fractions included Ib oligomer along with small amounts of Ia. **c,** Sample preparation for 2nd data. *top*, Density gradient ultracentrifugation. *bottom*, SDS-PAGE of fractions 17–20. Before 2nd data collection, Ia was added at three-fold molar excess.

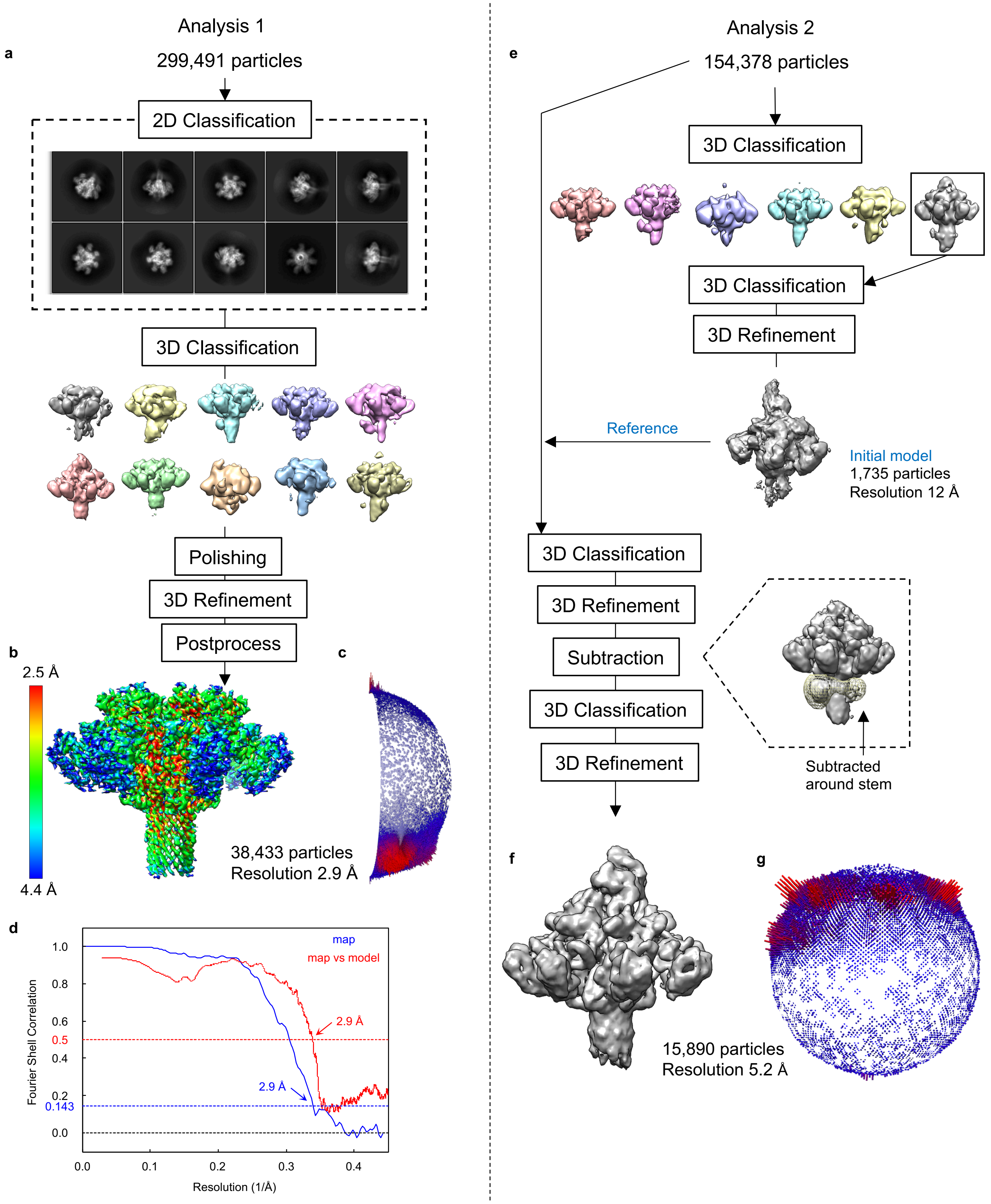


**Extended Data Figure 2 | Single particle analysis of 1st data.**
*left*, Ib-pore analysis. *right*, Ia-bound Ib-pore analysis. These analyses were performed individually from the same image set (1st data). **a,** Flow chart of cryo-EM image processing for Ib-pore. One of the 2D classes showed perfectly formed Ib-pore β-barrel although this class did not appear in subsequent 3D classes. **b,** Final 3D reconstruction map color-coded according to local resolution. **c,** Angular distribution of particles projected to the map. Angles of the projected particle is shown as 1/7 of a sphere because this analysis was performed with C7 symmetry. **d,** Gold-standard Fourier shell correlation (FSC) curve of final map and FSC curve for cross-validation between map and model. **e,** Flow chart of cryo-EM image processing for Ia-bound Ib-pore. After initial 3D refinement, density around the Ib pore stem was subtracted. **f,** Final 3D reconstruction map. **g,** Angular distribution of particles projected to the map.

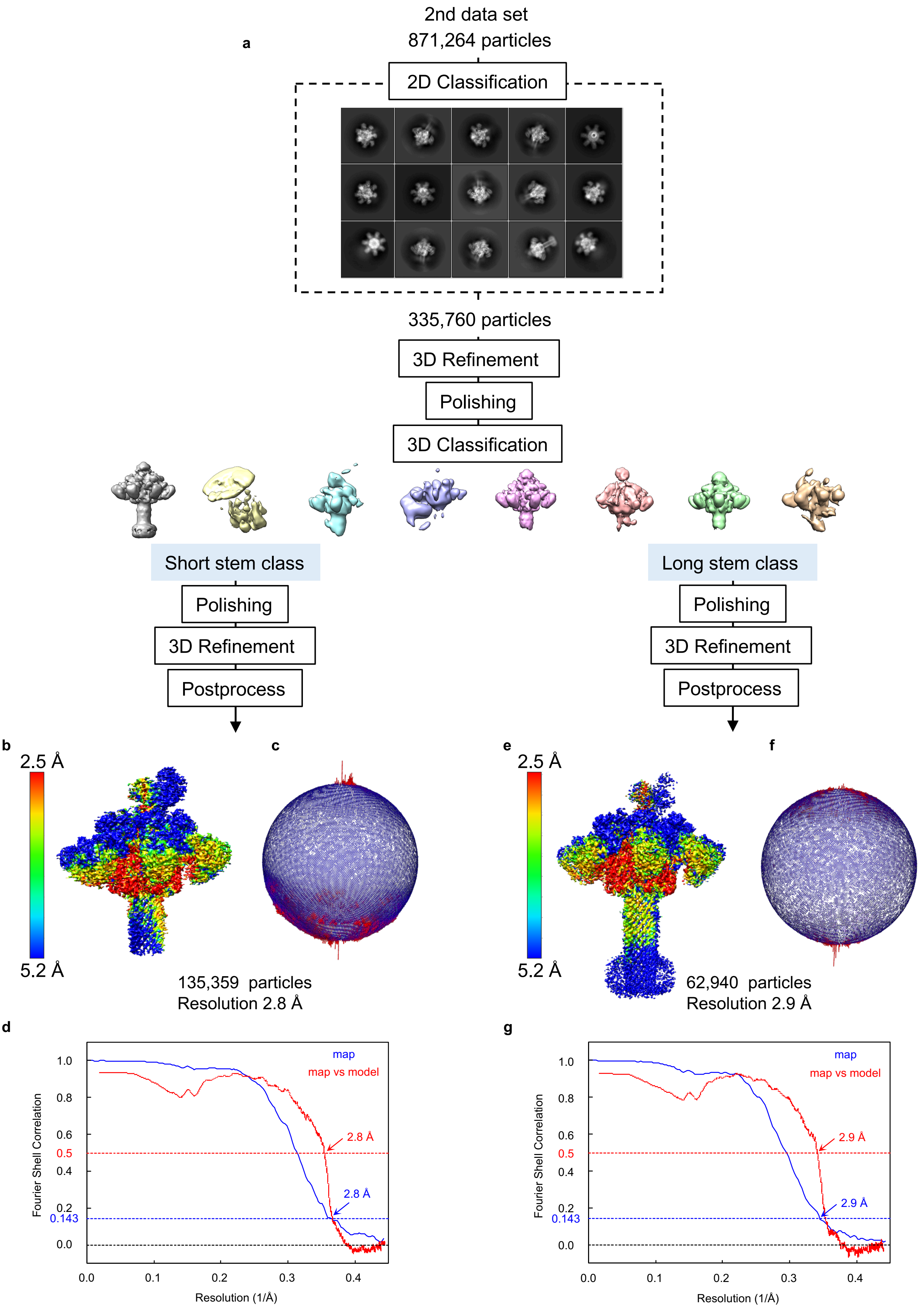
a
2nd data set
871,264 particles
2D Classification
335,760 particles
3D Refinement
Polishing
3D Classification
Short stem class
Polishing
3D Refinement
Postprocess
Long stem class
Polishing
3D Refinement
Postprocess
b
2.5 Å
5.2 Å
c
135,359 particles
Resolution 2.8 Å
e
2.5 Å
5.2 Å
f
62,940 particles
Resolution 2.9 Å
d
map
map vs model
2.8 Å
2.8 Å
Fourier Shell Correlation
Resolution (1/Å)
g
map
map vs model
2.9 Å
2.9 Å
Fourier Shell Correlation
Resolution (1/Å)

**Extended Data Figure 3 | Single particle analysis of 2nd data.**

**a,** Flow chart of cryo-EM image processing. **b,** Final 3D reconstruction map of Ia-bound Ib-pore with short stem, color-coded according to local resolution. Ia is clearly observed. **c,** Angular distribution of particles projected to the map shown in **b**. **d,** Gold-standard FSC curve of final map shown in **b** and FSC curve for cross-validation between the map and model. **e,** Final 3D reconstruction map of Ia-bound Ib-pore with long stem, color-coded according to local resolution. Ia is unclearer than map **b**. **f,** Angular distribution of particles projected to the map shown in **e**. **g,** Gold-standard FSC curve of final map shown in **e** and FSC curve for cross-validation between the map and model.

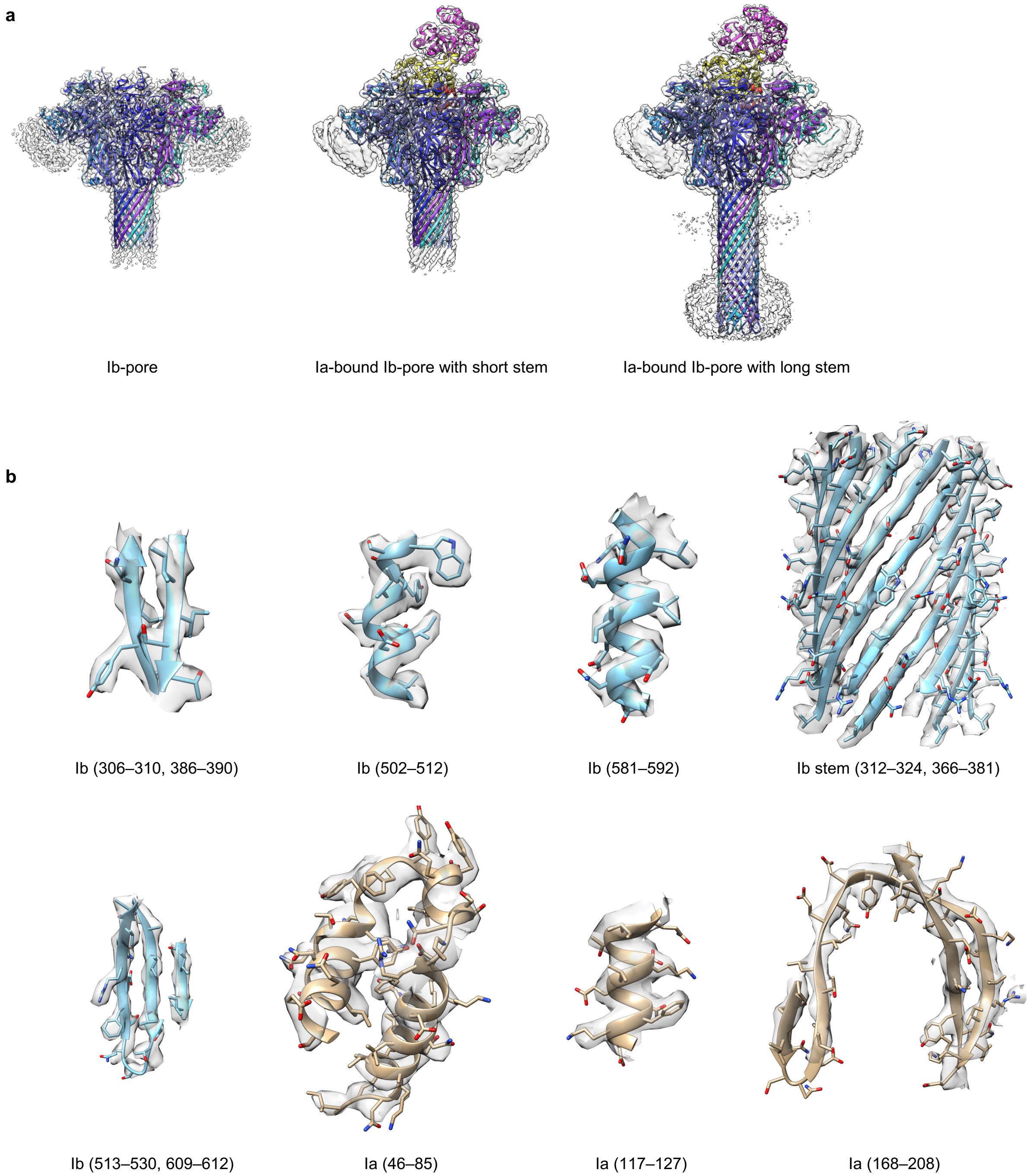


**Extended Data Figure 4 | Cryo-EM density maps and models.**
**a,** Overall maps and models. **b,** Representative cryo-EM density maps and models.

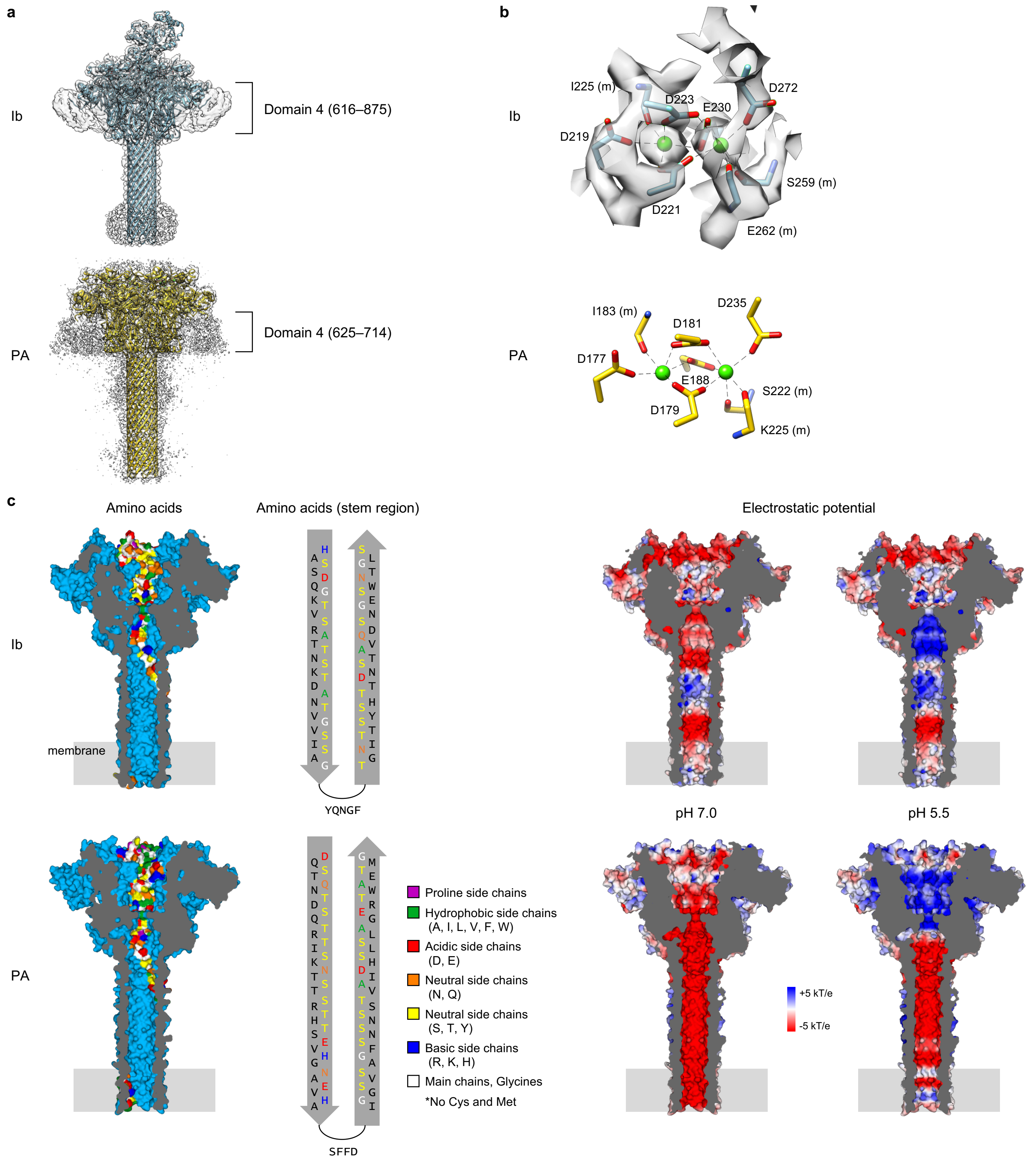


**Extended Data Figure 5 | Comparison between Ib and PA.**
**a,** Domain IV. Cryo-EM density maps and models are shown. PA PDB ID: 3J9C. Extra maps correspond to domain IV. The C-terminus of the Ib domain IV is invisible. **b,** Ca-binding sites. The calcium ions are shown as green spheres. Lowercase “m” in parentheses indicates that the main chain forms a coordination bond. **c,** Inner surfaces of pores. *left*, Colour coding of a protomer according to properties of amino acids. *right*, Cut-away surface electrostatic potential at pH 7.0 and 5.5.

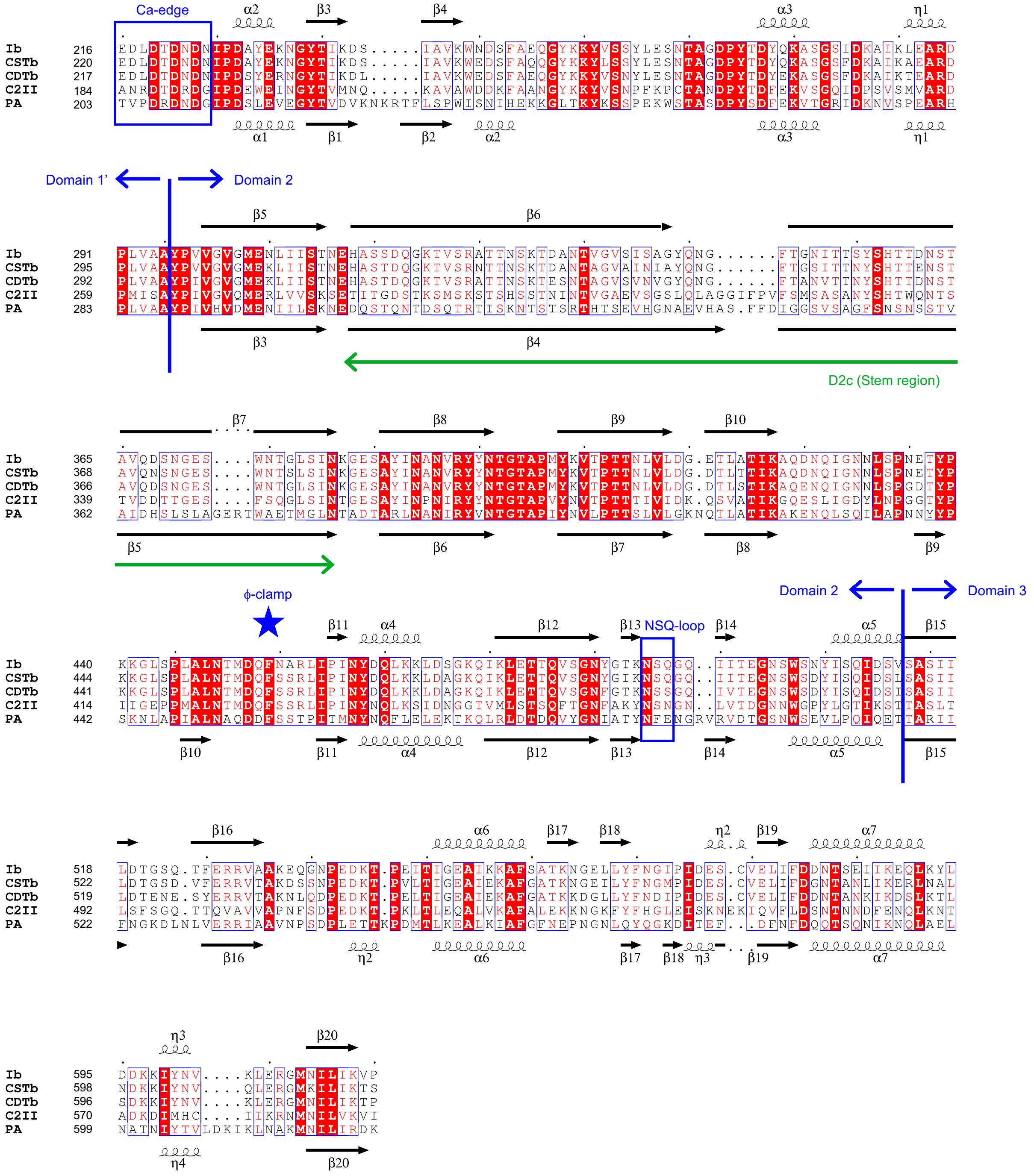


**Extended Data Figure 6 | Amino acid sequence alignment of domain 1′, 2, and 3.**
The secondary structures of Ib-pore and PA-pore (PDB ID: 3J9C) are shown above and below the sequence, respectively. The amino acids numbers including signal sequences are shown. Uniprot IDs: Ib, Q46221; CSTb, o06498; CDTb, o32739; C2II, o86171; PA, P13423.

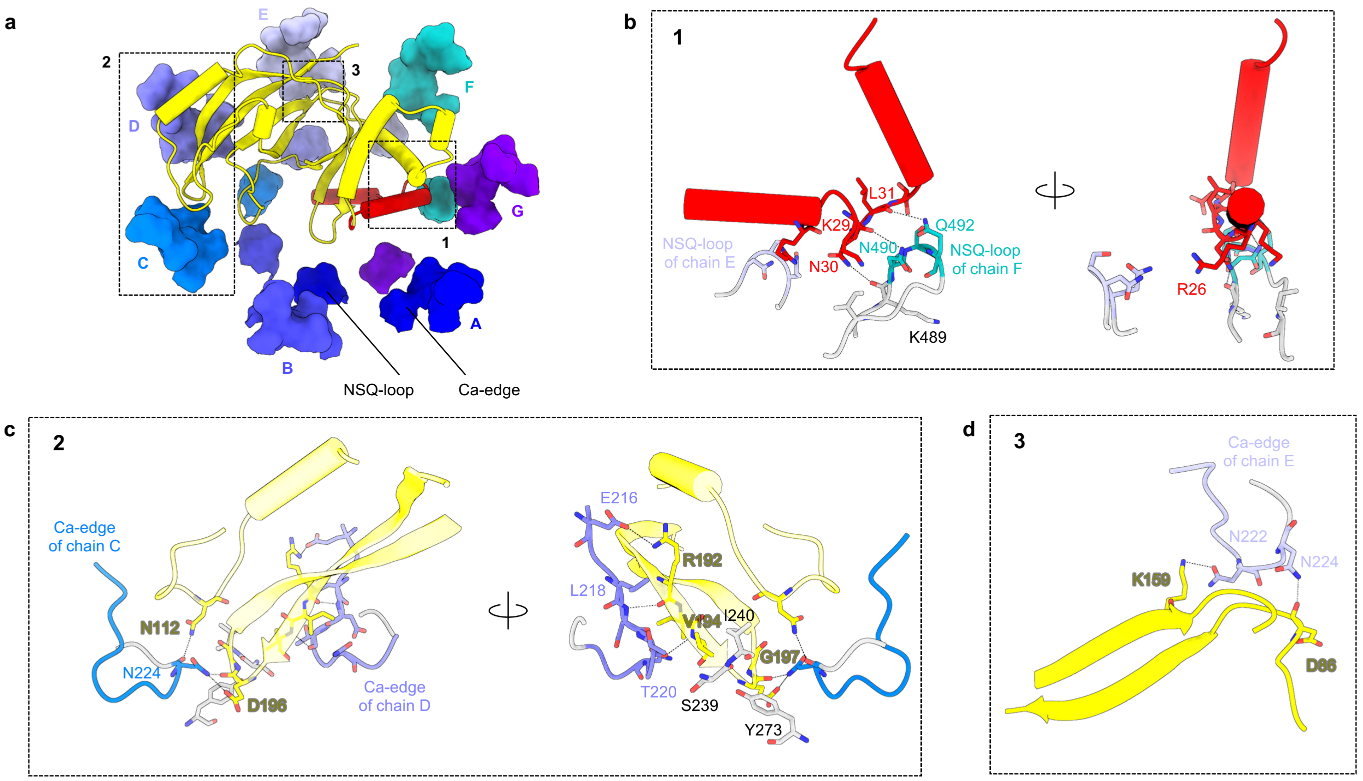


**Extended Data Figure 7 | Interactions between Ia and Ib.**

**a,** Overall view of Ia-Ib interactions. Ca-edges and NSQ-loops of Ib in surface model and N-terminal domain of Ia in cartoon model are shown. **b-d,** Close-up views of main interactions in three dash boxes 1–3.
