## Extended Table 1 for "Translocational unfolding in clostridial binary iota toxin complex"

**Cryo-EM data collection, refinement and validation statistics**

|  | Ib-pore  (EMDB-0721)  (PDB 6KLX) | Ia-bound Ib-pore with short stem  (EMDB-0713)  (PDB 6KLO) | Ia-bound Ib-pore with long stem  (EMDB-0720)  (PDB 6KLW) |
| --- | --- | --- | --- |
| **Data collection and processing** |  |  |  |
| Magnification | ×59,000 | ×59,000 | ×59,000 |
| Voltage (kV) | 300 | 300 | 300 |
| Electron exposure (e–/Å^2^) | 50 | 50 | 50 |
| Defocus range (μm) | -0.8 to -2.5 | -0.8 to -2.5 | -0.8 to -2.5 |
| Pixel size (Å) | 1.13 | 1.13 | 1.13 |
| Symmetry imposed | C7 | C1 | C1 |
| Initial particle images (no.) | 299,491 | 871,264 | 871,264 |
| Final particle images (no.) | 38,433 | 135,359 | 62,940 |
| Map resolution (Å)  FSC threshold | 2.9  0.143 | 2.8  0.143 | 2.9  0.143 |
| Map resolution range (Å) | 2.5–4.4 | 2.5–5.2 | 2.5–5.2 |
| **Refinement** |  |  |  |
| Initial model used (PDB code) | 3J9C | 3J9C, 1GIQ | 3J9C, 1GIQ |
| Model resolution (Å)  FSC threshold | 2.9  0.5 | 2.8  0.5 | 2.9  0.5 |
| Model resolution range (Å) |  |  |  |
| Map sharpening *B* factor (Å^2^) |  |  |  |
| Model composition  Non-hydrogen atoms  Protein residues  Ligands | 20055  2576  14 Ca^2+^ | 23275  2972  14 Ca^2+^ | 25179  3238  14 Ca^2+^ |
| *B* factors (Å^2^)  Protein  Ligand | 49.67  40.87 | 57.47  37.59 | 51.02  27.50 |
| R.m.s. deviations  Bond lengths (Å)  Bond angles (°) | 0.008  0.686 | 0.005  0.591 | 0.005  0.653 |
| Validation  MolProbity score  Clashscore  Poor rotamers (%) | 1.83  7.01  0.00 | 1.72  5.93  0.00 | 1.80  6.84  0.00 |
| Ramachandran plot  Favored (%)  Allowed (%)  Disallowed (%) | 93.21  6.79  0.00 | 94.05  5.95  0.00 | 93.51  6.49  0.00 |
